# Down syndrome postmortem brains exhibit reduction in [^18^F]nifene binding to α4β2* nicotinic acetylcholinergic receptors

**DOI:** 10.64898/2026.09.09.750420

**Authors:** Fariha Karim, Nancy Paola Chavez-Jacobo, Christopher Liang, Elizabeth Head, Jogeshwar Mukherjee

## Abstract

Alzheimer’s disease (AD) pathology including amyloid beta (Aβ) plaques and tau tangles accumulate with age in Down Syndrome (DS). Cholinergic abnormalities such as expression of nicotinic acetylcholine receptors (nAChRs) adversely contribute to cognitive decline and neurodegeneration in AD with similar features possible in DSAD. As a radiotracer for α4β2* nAChRs, [^18^F]nifene was quantitatively evaluated in the frontal cortex (FCX) and temporal cortex (TCX) of DSAD, AD, and cognitively normal (CN) brain tissue using autoradiography. Anti-tau and anti-Aβ immunostaining in adjacent sections confirmed the presence of tau tangles and Aβ plaques. [^18^F]Nifene binding in brain sections demonstrated significantly more binding in gray matter (GM) than white matter (WM), with FCX exhibiting more binding than TCX (OptiQuant). Nicotine substantially displaced [^18^F]nifene binding in adjacent sections, resulting in average GM/nicotine ratios (DSAD=9.74, AD=10.5, CN=17; suggesting a 38% decrease in AD and a 43% decrease in DSAD) and WM/nicotine ratios (DSAD=2.73, AD=3.31, CN=5.62; suggesting a 41% decrease in AD and a 51% decrease in DSAD). [^18^F]Nifene was correlated with [^125^I]IBETA (Aβ), showing a strong positive relationship between α4β2* nAChRs and Aβ in DSAD. [^18^F]Nifene GM/nicotine and WM/nicotine ratios were positively correlated with age in TCX and FCX. The findings of this study suggest that [^18^F]nifene has potential use in diagnostic investigations of the cholinergic system in DSAD using positron emission tomography. The study also points to a possible need for earlier therapeutic interventions to address the α4β2* nAChRs deficit in DSAD.

**SIGNIFICANCE STATEMENT:** People with Down syndrome are vulnerable to Alzheimer disease along with decreased cholinergic function, which may be linked to cognitive decline. The accumulation of amyloid plaques and tau associated with Alzheimer’s disease initiation and progression may be linked to cholinergic dysfunction in Down syndrome. Here we report decreases in α4β2 nicotinic acetylcholine receptors in Down syndrome frontal and temporal cortex. This receptor subtype is important for several brain functions, and loss of this receptor therefore contributes to some of the cognitive deficits seen in Down syndrome with increasing age.

## 1. INTRODUCTION

Most, if not all, individuals with Down syndrome (DS) develop Alzheimer’s disease (AD) neuropathology including amyloid beta (Aβ) plaques and neurofibrillary tangles (Rafii et al. 2025). The triplication of chromosome 21 includes overexpression of the amyloid precursor protein (APP) gene, that is thought to drive widespread age associated amyloid deposition (Wisniewski et al. 1985; Doran et al. 2017; Ovchinnikox et al. 2018). Accumulation of Aβ plaques in DS appears to begin in the striatum, followed rapidly (∼3 years) by neurofibrillary tangles that start in the entorhinal cortex and spread to the temporal cortex (Annus et al. 2016; Braak & Braak 1991; Cohen et al. 2018; Zammit et al. 2023). This accumulation of AD pathology and rapid spread is associated with dementia and cognitive decline (Annus et al. 2016).

The development of AD pathology is thought to arise with the development of cholinergic dysfunction (Hampel et al. 2018). The cholinergic system plays a crucial role in memory, learning, and several brain functions required for brain homeostasis and plasticity. Cholinergic basal forebrain deficits are associated with poor cognitive function and performance in adults with AD and older healthy adults (Richter et al. 2022). The basal forebrain is also susceptible to neurofibrillary tangles and Aβ deposition, which correlate with cholinergic neuron loss (Hampel et al. 2018). This susceptibility may be a critical link between cognitive impairment and neurodegeneration in DS and AD (Martinez et al. 2021). Morphological similarities between DS and AD include the presence of senile plaques and neurofibrillary tangles within the cerebral cortex and hippocampus, progressively deteriorating cholinergic nerve terminal function (Mann et al. 1985; Isacson et al. 2002). In addition to the overexpression of APP possibly contributing to cholinergic system decline and development of AD pathology, other genes on chromosome 21 may also contribute (Russell et al. 2024). During the development of AD in DS, cholinergic basal forebrain volume decreases with age as measured by MRI and is strongly correlated with amyloid and tau changes measured by PET (Rozalem Aranha et al. 2023). Basal forebrain atrophy may indicate AD-related cholinergic neurodegeneration in DS.

Mechanisms of cholinergic neurodegeneration can be traced to the nicotinic acetylcholine receptors (nAChR) that support cognitive function. The most abundant subtype, α4β2 nAChRs, reside in brain regions responsible for cognition, which is impaired in AD (Lombardo & Maskos 2014; Sabri et al. 2018). Changes in cholinergic receptor expression are apparent in DS such as elevation of the β2 subunit (Engidawork et al. 2001 Isacson et al. 2002). The vulnerability of basal forebrain cholinergic neurons (BFCN) to neurodegeneration, vital for cholinergic innervation regulating attention and memory, suggests a link to cognitive decline in DS and AD (Martinez et al. 2021).

PET studies may detect cholinergic abnormalities using non-invasive and sensitive imaging. [^18^F]-fluoroethoxybenzovesamicol ([^18^F]FEOBV) is a radiotracer that binds to vesicular acetylcholine transporter in cholinergic nerve terminals (Mulholland et al. 1998). As one of the first radiotracers used to quantify cholinergic deficits, [^18^F]FEOBV demonstrates reductions in vesicular acetylcholine transporters that are significantly associated with cholinergic and cognitive decline in AD (Aghourian et al. 2017). In DS, cholinergic terminal density is initially elevated in early adulthood before a steep decline, disrupting cholinergic signaling measured by [^18^F]FEOBV PET (Russell et al. 2025). Paradoxically, [^18^F]FEOBV uptake throughout the brain appears to be higher in adults with DS but there is a stronger negative correlation with age when compared with similarly aged neurotypical adults (Russell et al. 2025).

Significant effort has been made to study two prominent brain nicotinic acetylcholinergic receptors (nAChRs), namely the heteromeric α4β2 and the homomeric α7 nAChRs and their role in neurodegeneration. However, their role in DSAD pathophysiology has not been described but several in vivo PET imaging and postmortem studies have been reported in AD and Parkinson’s disease (PD). Using different radioligands and imaging methods, results have been mixed. The α7 nAChRs that are abundant in brain regions implicated in learning and memory, such as the hippocampus and cortex, have not been observed to be affected in AD and PD (Ngo et al. 2025, Karim et al. 2025,). The α4β2 nAChRs, which have higher affinity for acetylcholine and are abundant in the thalamus, cortical brain regions including subiculum and hippocampus, are significantly reduced in AD (Karim et al. 2025). In PD, observations of α4β2 nAChRs are mixed, but a recent report using [^18^F]nifene as the radioligand suggests a significant increase in binding in subiculum and hippocampus of postmortem PD cases (Mukherjee et al. 2026). It may also be noted that both, heteromeric α4β2 and the homomeric α7 nAChRs are ion-channels. Therefore, diagnostic agents for ion-channel associated proteins pose additional challenges in the evaluation of their role in health and disease (Shah et al. 2024).

PET imaging of α4β2* nAChRs using [^18^F]nifene shows wide-spread distribution of this receptor in the human brain and provides valuable insight into the intact cholinergic pathways throughout the brain in a normal subject (Figure 1) (Mukherjee et al. 2018). In addition to the grey matter (GM) regions, white matter (WM) has also been an area of interest for [^18^F]nifene binding both in small animals and humans (Bieszczad et al. 2012; Mukherjee et al. 2018). Thalamocortical pathways containing α4β2* nAChRs can influence learning and memory tasks in rodents (Bieszczad et al. 2012). White matter thalamic radiations containing α4β2* nAChRs can play a critical role in brain networks and should be evaluated in neurodegeneration (Mukherjee et al. 2018).

**Figure 1:**
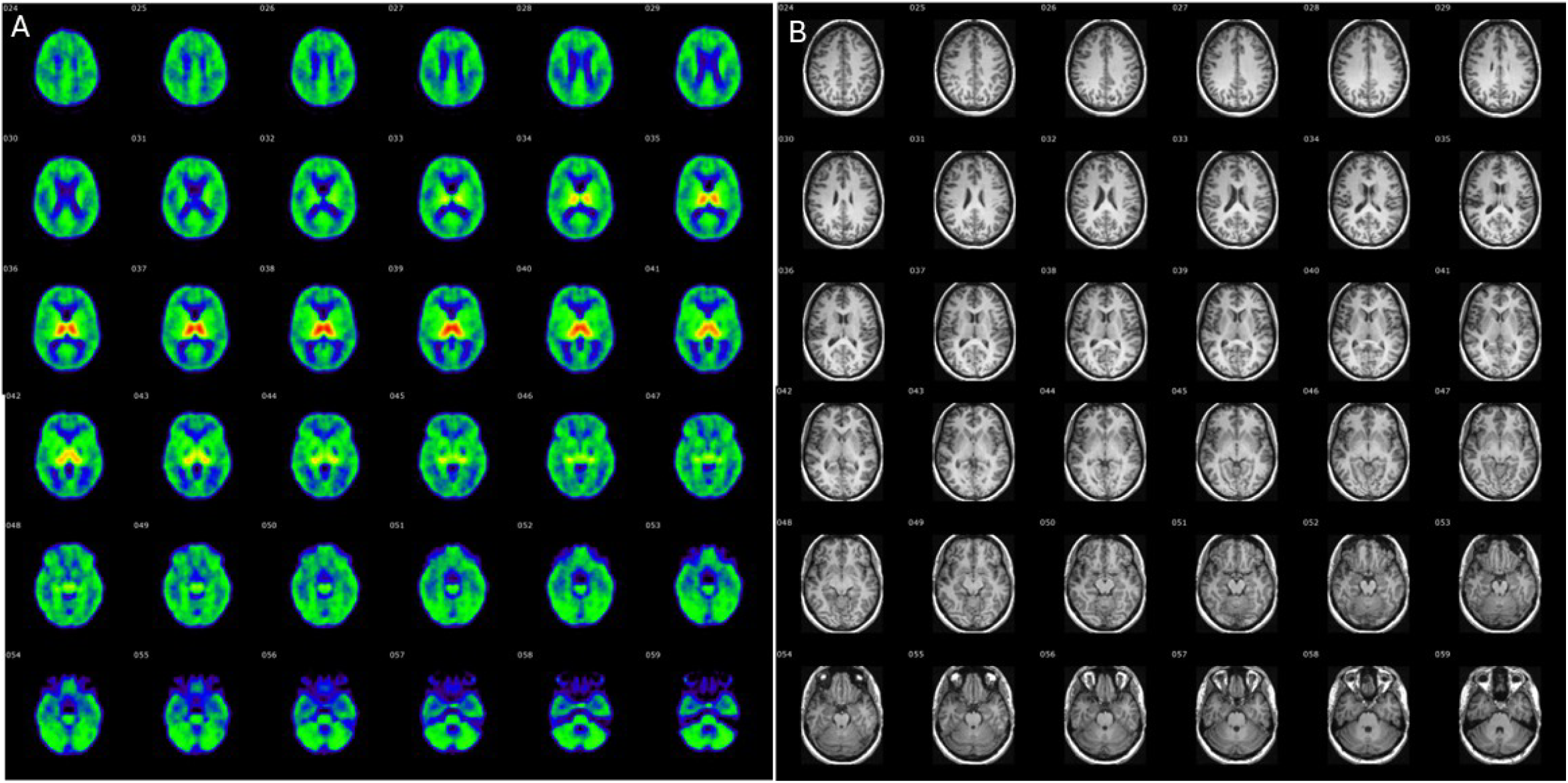
[^18^F]Nifene for imaging human α4β2* nAChRs: (A). Transaxial PET images of [^18^F]nifene binding to cognitively normal subject from dorsal to ventral planes (Mukherjee et al. 2018). Thalamus shows the highest [^18^F]nifene binding in red. The wide-spread distribution of α4β2* nAChRs in the human cortical and sub-cortical structures is evident. (B). Corresponding transaxial MRI images of the same subject showing grey matter and white matter in the same brain slices.

Our recent postmortem studies in DSAD shows extensive Aβ and tau pathology comparable to AD postmortem brains (Biju et al. 2026; Karim et al. 2026). Our recent postmortem study shows lower levels of [^18^F]nifene binding in the hippocampus and subiculum in AD brains. We proposed that a similar reduction may be expected in DSAD, due to the presence of similar neuropathology. Therefore, we have tested [^18^F]nifene binding in DSAD in the frontal cortex (FCX), as an area affected early by Aβ and temporal cortex (TCX), which exhibits early tau accumulation (Lemoine et al. 2020; Rafii et al. 2025). This study aims to investigate [^18^F]nifene binding in DSAD, AD, and cognitively normal (CN) FCX and TCX cases positive for Aβand tau for insight into proteinopathy and age relationships. The study also investigated [^18^F]nifene binding in the WM of DSAD in comparison to AD and CN.

## 2. MATERIALS AND METHODS

### 2.1 Radiopharmaceuticals

#### [^18^F]Nifene

The radiosynthesis of [^18^F]nifene was performed using nucleophilic displacement of the nitro group in *N*-BOC-nitronifene precursor (prepared in-house, Pichika et al. 2006) by [^18^F]fluoride (PETNET, Inc.) in an automated synthesizer followed by deprotection using previously described procedures (Pichika et al. 2006; Campoy et al. 2021; Bhuiyan et al. 2024). [^18^F]nifene radiochemical purity was >98%, chemical purity was >95%, and the molar activity was measured to be >70 GBq/µmol (>2 Ci/µmol) at the end of synthesis.

#### [^125^I]IBETA

[^125^I]IBETA is a radiotracer for imaging Aβ plaques and exhibited selective binding to human AD brain Aβ plaques (Nguyen et al. 2022; Mondal et al. 2023). The radiosynthesis of [^125^I]IBETA used in-house precursor (Nguyen et al. 2022) and was synthesized in >95% purity with a measured molar activity >70 GBq/µmol (>2 Ci/µmol) at the end of synthesis.

#### [^18^F]MK-6240

[^18^F]MK-6240 is a radiotracer for imaging Tau (Walji et al. 2016; Karim et al. 2026). The radiosynthesis of [^18^F]MK-6240 was performed with [^18^F]fluoride using previously described procedures (Karim et al. 2026). Radiochemical purity of [^18^F]MK-6240 was >98% and chemical purity was found to be >95% (molar activity >70 GBq/µmol (>2 Ci/µmol) at the end of synthesis.

### 2.2 Postmortem Human Brain

Human postmortem brain tissue samples of CN, AD, and DSAD cases (n=5), each including FCX and TCX, were obtained from the UCI Alzheimer disease research center at UCI Memory Impairment and Neurological Disorders (MIND) institute. Case demographics are provided in Table 1. All AD and DSAD cases were positive for tau and Aβ. Brain slices, 10 µm thick on Fisher glass slides, were collected using a Leica 1850 cryotome set to -20 °C. Each slide contains 2-3 brain sections. Typical sizes of the collected brain sections ranged from approximately 1 to 1.5 cm^2^. All slides were then stored at -80°C. All postmortem human brain studies were approved by the Institutional Biosafety Committee of University of California, Irvine.

**Table 1.**
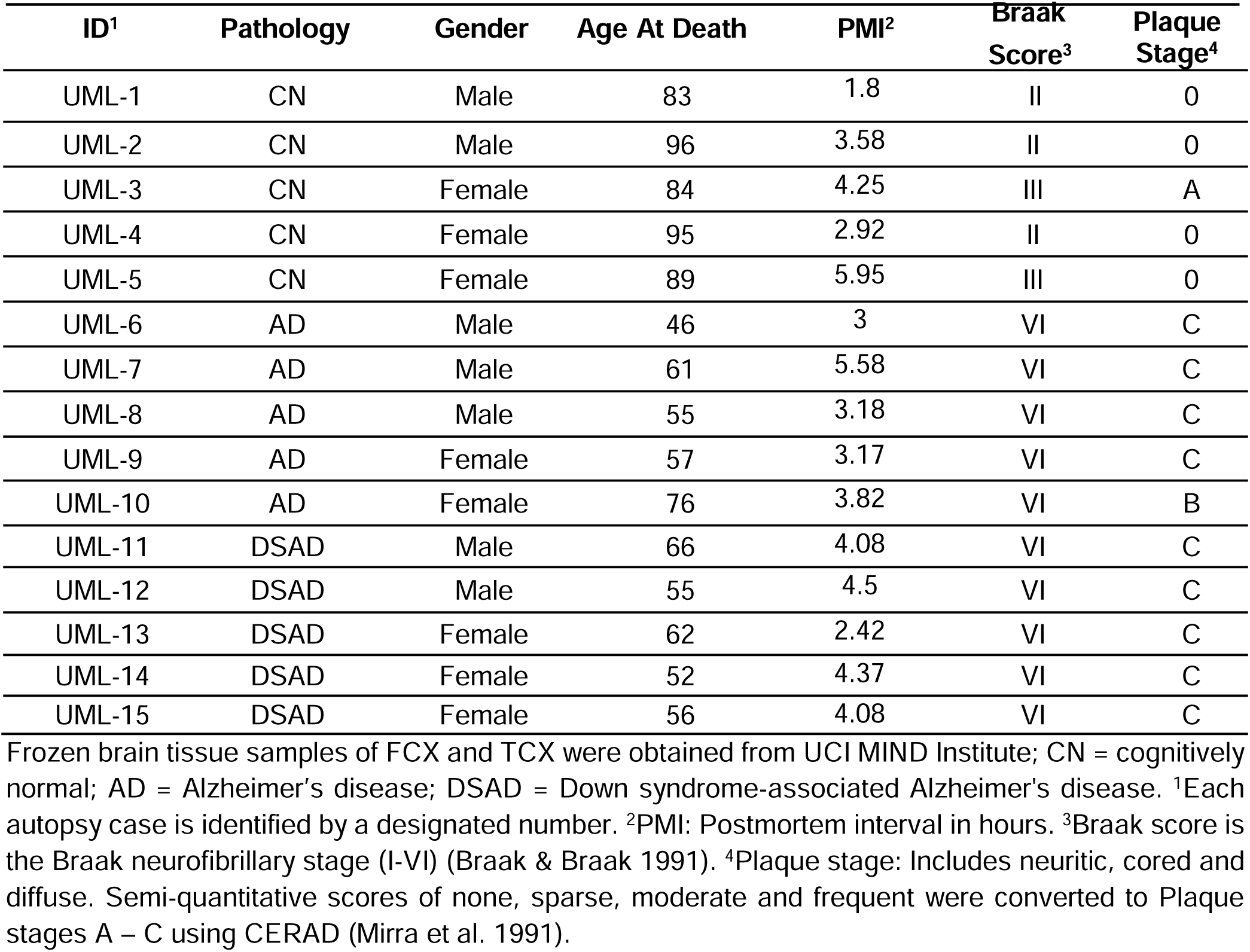
Patient Case samples and data.

| ID<sup>1</sup> | Pathology | Gender | Age At Death | PMI<sup>2</sup> | Braak Score<sup>3</sup> | Plaque Stage<sup>4</sup> |
| --- | --- | --- | --- | --- | --- | --- |
| UML-1 | CN | Male | 83 | 1.8 | II | 0 |
| UML-2 | CN | Male | 96 | 3.58 | II | 0 |
| UML-3 | CN | Female | 84 | 4.25 | III | A |
| UML-4 | CN | Female | 95 | 2.92 | II | 0 |
| UML-5 | CN | Female | 89 | 5.95 | III | 0 |
| UML-6 | AD | Male | 46 | 3 | VI | C |
| UML-7 | AD | Male | 61 | 5.58 | VI | C |
| UML-8 | AD | Male | 55 | 3.18 | VI | C |
| UML-9 | AD | Female | 57 | 3.17 | VI | C |
| UML-10 | AD | Female | 76 | 3.82 | VI | B |
| UML-11 | DSAD | Male | 66 | 4.08 | VI | C |
| UML-12 | DSAD | Male | 55 | 4.5 | VI | C |
| UML-13 | DSAD | Female | 62 | 2.42 | VI | C |
| UML-14 | DSAD | Female | 52 | 4.37 | VI | C |
| UML-15 | DSAD | Female | 56 | 4.08 | VI | C |

Frozen brain tissue samples of FCX and TCX were obtained from UCI MIND Institute; CN = cognitively normal; AD = Alzheimer's disease; DSAD = Down syndrome-associated Alzheimer's disease. <sup>1</sup>Each autopsy case is identified by a designated number. <sup>2</sup>PMI: Postmortem interval in hours. <sup>3</sup>Braak score is the Braak neurofibrillary stage (I-VI) (Braak & Braak 1991). <sup>4</sup>Plaque stage: Includes neuritic, cored and diffuse. Semi-quantitative scores of none, sparse, moderate and frequent were converted to Plaque stages A – C using CERAD (Mirra et al. 1991).

### 2.3 [^18^F]Nifene Autoradiography for α4β2* nAChR

Slides of brain slices were inserted into glass chambers to warm to room temperature and preincubate in Tris buffer (50 mmol/L Tris, 120 mmol/L NaCl, 5 mmol/L KCl, 2.5 mmol/L CaCl_2_, 1 mmol/L MgCl_2_, pH 7.4) for 15 min. After discarding the preincubation buffer, [^18^F]nifene in Tris buffer pH 7.4 (60 mL; 37 kBq/mL) was poured to each chamber. All chambers were incubated at 25°C for 1 hr. Nonspecific binding of [^18^F]nifene was measured of adjacent brain slices in the presence of 300 |jM nicotine. The incubation buffer was disposed, and the slides were washed as follows: cold Tris buffer twice for 3 minute each time and cold deionized water for 1 minute. The slides were air dried and apposed to phosphor films (Perkin Elmer Multisensitive, Medium MS) inside film cassettes overnight. The next day, films were read using the Cyclone Phosphor Imaging System (Packard Instruments Co).

Autoradiography of adjacent brain slices for Aβ Plaques and Tau using [^125^I]IBETA and [^18^F]MK-6240 respectively followed previously reported methods (Biju et al. 2026; Karim et al. 2026).

### 2.4 Optiquant Image Analysis

Regions of interest (ROIs) in the GM and WM were drawn on the autoradiographic images of FCX and TCX brain slices using the analysis program Optiquant (Packard Instruments Co, version 5.0). In each ROI, pixels of the autoradiographic image measured radiotracer binding in digital light units (DLU/mm^2^). Background activity levels were subtracted from all ROIs, resulting in net binding. Higher DLU/mm^2^ from autoradiography indicated higher [^18^F]nifene, [^125^I]IBETA, and [^18^F]MK-6240 binding.

### 2.5 Immunohistochemistry

Adjacent brain slices of all cases were immunostained for tau and Aβ plaques by UCI Pathology core services using Ventana BenchMark Ultra protocols. DAKO polyclonal antibody detects all 6 six isoforms of tau and was used for total tau at a dilution 1:3000, A0024 (Agilent, CA, USA) using reported protocols (Ercan et al. 2017). An additional set of adjacent brain slices were immunostained with anti-Aβ Biolegend 803015 (Biolegend, CA, USA) which is reactive to amino acid residue 1-16 of β-amyloid. The Ventana Roche slide scanner was used to capture images of the anti-tau and anti-Aβ immunostained slides to visualize in QuPath (version 4.4).

### 2.6 Statistical analysis

Microsoft Excel 16 and GraphPad Prism 10 were used for all statistical analysis. Parametric student’s t-tests and one-way ANOVA with Dunnett’s multiple comparisons test determined statistical power (p-value <0.05). Error bars symbolize mean ± standard deviation within each group. Pearson’s correlation assesses biomarker relationships and aging effects. Simple linear regressions obtain a R^2^ value that indicates the linearity of the relationship. The ROUT method of identifying outliers (Q=1%) was performed in GraphPad Prism 10 and identified no outliers.

## 3. RESULTS

### 3.1 DSAD Postmortem Human FCX and TCX

Figure 2 summarizes [^18^F]nifene binding in DSAD FCX (Figure 2A-E) and TCX (Figure 2F-J) sections using a representative case (13). All cases were positive for tau and Aβ in anti-tau (Figure 2B,G) and anti-Aβ (Figure 2C,H) immunostaining of adjacent slices. Autoradiographic images show total [^18^F]nifene binding in FCX (Figure 2D) and TCX (Figure 2I) sections. Nicotine (300 |jM) successfully displaced [^18^F]nifene in all DSAD cases from both GM and WM regions (Figure 2E,J). [^18^F]Nifene binding was significantly greater in GM than WM across all the cases in both brain regions (Figure 2K). Significant average ratios between TCX GM/TCX WM= 3.18 and FCX GM/FCX WM= 3.12 were measured, suggesting similar levels of [^18^F]nifene binding in the two brain regions.

**Figure 2:**
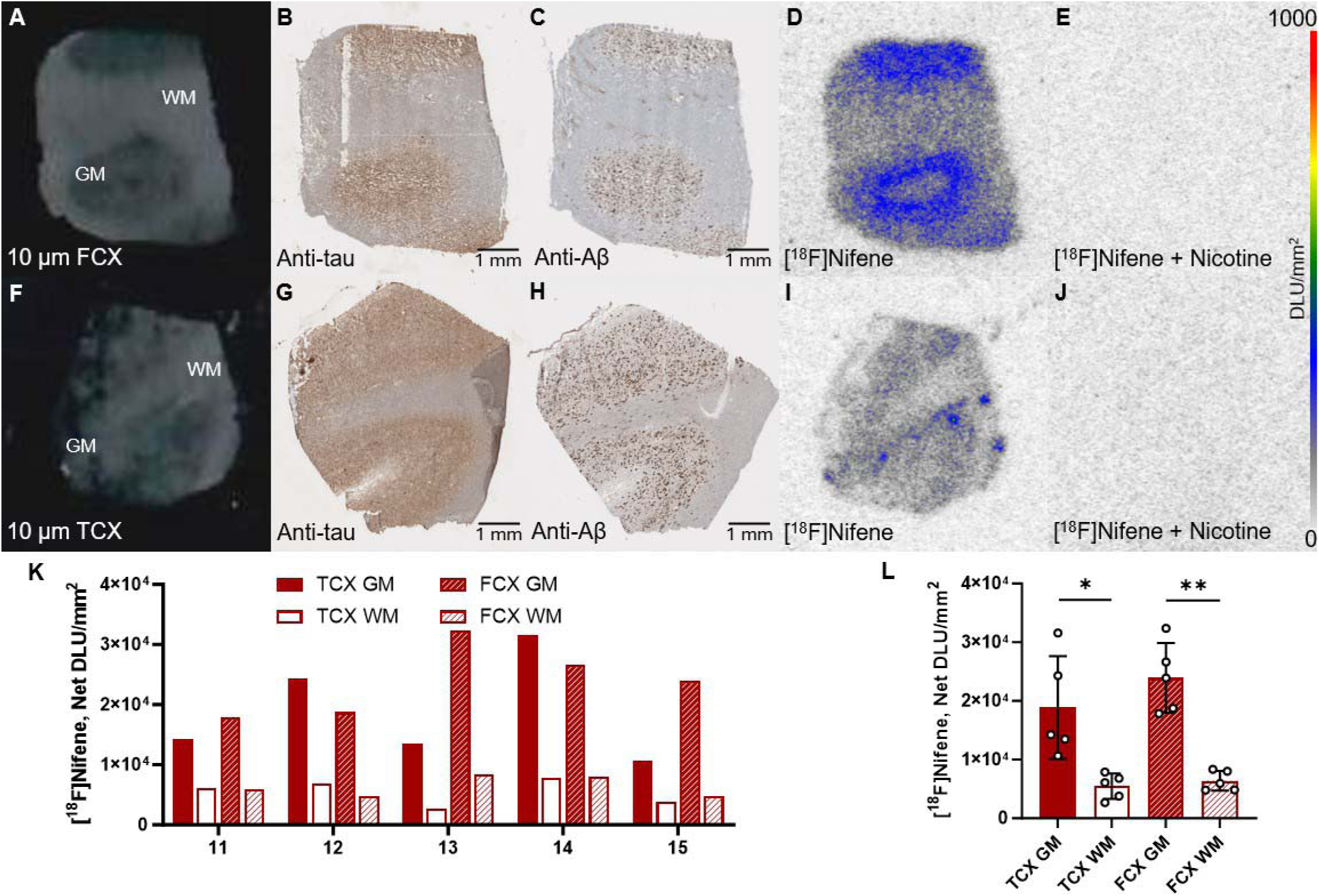
DSAD cases [^18^F]nifene: (A-E). DSAD FCX case 13; (F-J) DSAD TCX case 13; (A, E). DSAD postmortem human brain slice with labeled GM and WM; (B, G). DSAD anti-tau immunostain (1 mm); (C, H). DSAD anti-Aβ immunostain (1 mm); (D, I). Total [^18^F]nifene binding to DSAD (autoradiography scale bar: 0-1000 DLU/mm^2^); (E, J). [^18^F]nifene binding to adjacent brain section in competition with nicotine (300 |jM) (autoradiography scale bar: 0-1000 DLU/mm^2^); (K). [^18^F]nifene binding in DLU/mm^2^ to GM and WM of all DSAD TCX and FCX; (L). Average [^18^F]nifene binding in DLU/mm^2^ to GM and WM within DSAD TCX and FCX. Paired two-tailed parametric t-tests determined statistical significance between GM and WM (*p<0.05,***p<0.001).

### 3.2 AD Postmortem Human FCX and TCX

Similarly, AD FCX and TCX sections were studied [^18^F]nifene in AD in order to compare with similar brain regions of DSAD (Figure 3). Positivity for tau and Aβ was verified in all cases and in anti-tau immunostaining (Figure 3B,G). Total binding of [^18^F]nifene (Figure 3D,I) was evaluated alongside [^18^F]nifene binding being displaced by nicotine (Figure 3E,J). The extent of [^18^F]nifene binding was significantly greater in GM than WM in all AD cases for TCX and FCX (Figure 3K-L). Like DSAD cases, [^18^F]nifene binding was significantly greater in GM than WM across all the AD cases in both brain regions (Figure 2K). Average ratios between TCX GM/TCX WM= 3.57 and FCX GM/FCX WM= 2.81 were measured, suggesting some differences of [^18^F]nifene binding from the DSAD cases. It should be noted that the variability in WM [^18^F]nifene binding had a direct effect on the variability of these ratios.

**Figure 3:**
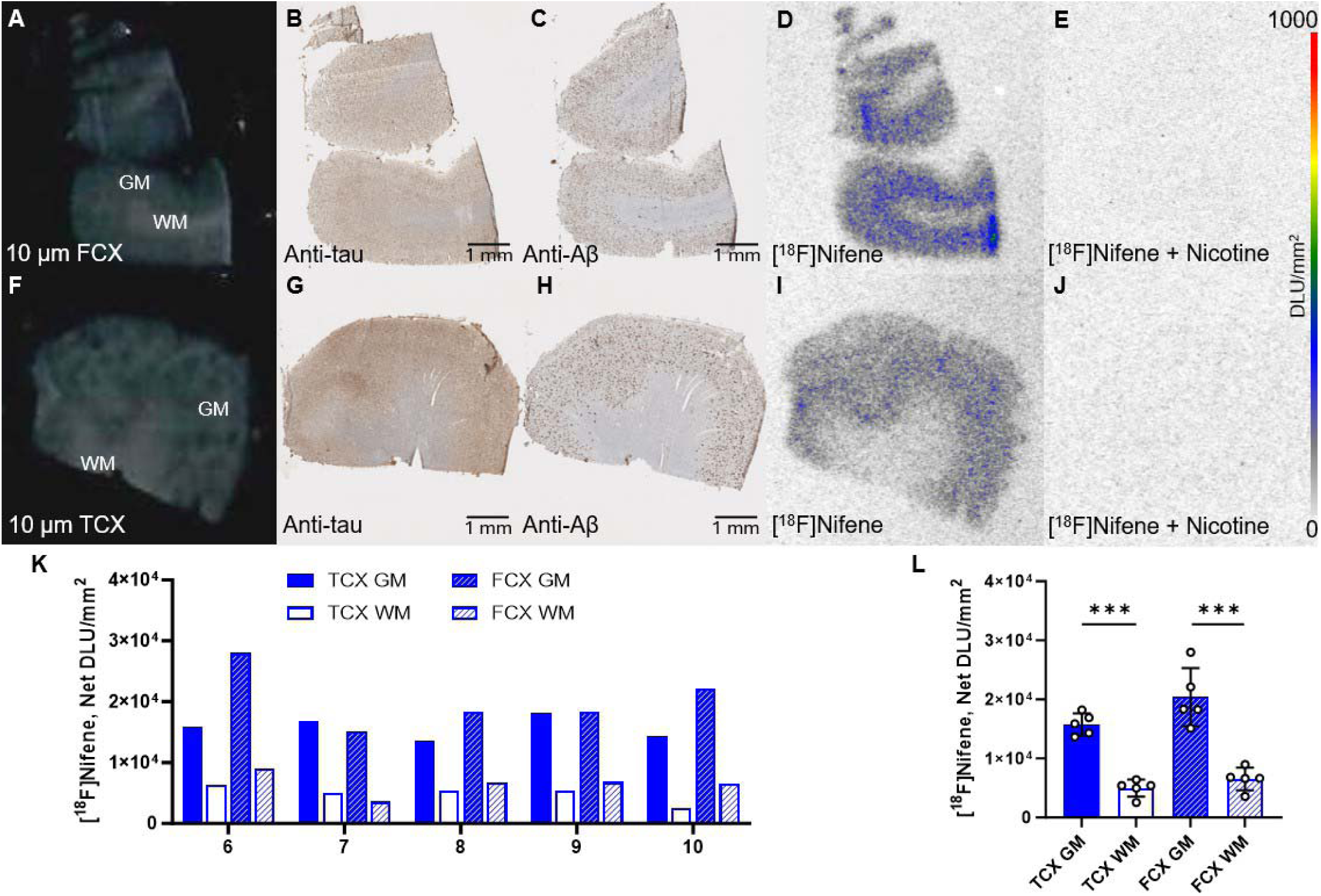
AD cases [^18^F]nifene: (A-E). AD FCX case 10; (F-J) AD TCX case 10; (A, E). AD postmortem human brain slice with labeled GM and WM; (B, G). AD anti-tau immunostain (1 mm); (C, H). AD anti-Aβ immunostain (1 mm); (D, I). Total [^18^F]nifene binding to AD (autoradiography scale bar: 0-1000 DLU/mm^2^); (E, J). [^18^F]nifene binding to adjacent brain section in competition with nicotine (300 |jM) (autoradiography scale bar: 0-1000 DLU/mm^2^); (K). [^18^F]nifene binding in DLU/mm^2^ to GM and WM of all AD TCX and FCX; (L). Average [^18^F]nifene binding in DLU/mm^2^ to GM and WM within AD TCX and FCX. Paired two-tailed parametric t-tests determined statistical significance between GM and WM (*p<0.05,***p<0.001).

### 3.3 CN Postmortem Human FCX and TCX

[^18^F]Nifene binding was evaluated in all CN cases in FCX and TCX (Figure 4). The localization of total [^18^F]nifene binding is provided from autoradiographic images for both FCX (Figure 4B) and TCX (Figure 4E) brain slices. To resolve potential nonspecific binding, [^18^F]nifene binding was competed with 300 |jM nicotine (Figure 4C,F). Quantification of total [^18^F]nifene binding shows a significant difference between GM and WM among all cases in both TCX and FCX sections (Figure 4G-H). In the CN cases, ratios between TCX GM/TCX WM= 3.46 and FCX GM/FCX WM= 3.74 were measured.

**Figure 4:**
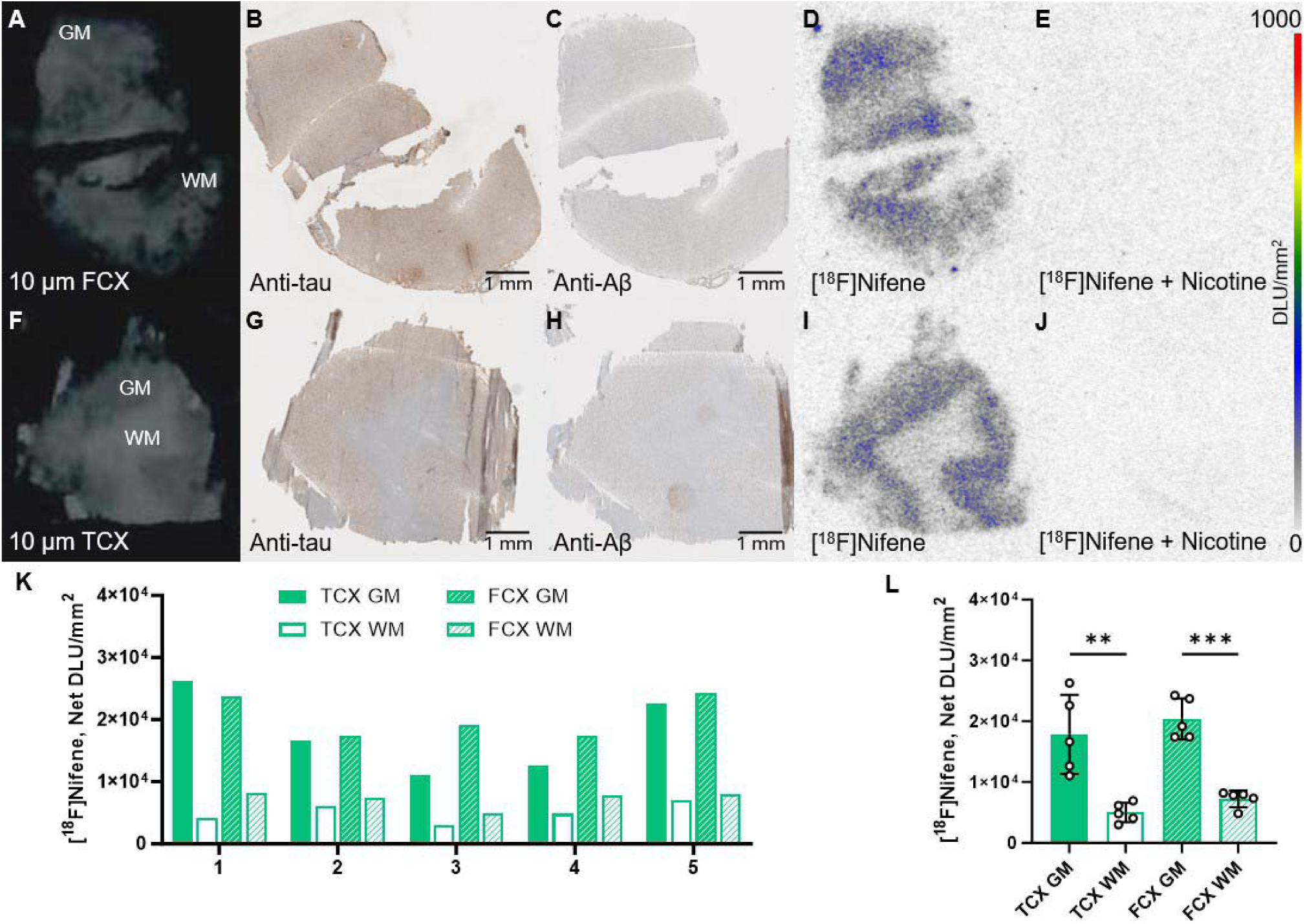
CN cases [^18^F]nifene: (A-E). CN FCX case 5; (F-J) CN TCX case 5; (A, E). CN postmortem human brain slice with labeled GM and WM; (B, G). CN anti-tau immunostain (1 mm); (C, H). CN anti-Aβ immunostain (1 mm); (D, I). Total [^18^F]nifene binding to CN (autoradiography scale bar: 0-1000 DLU/mm^2^); (E, J). [^18^F]nifene binding to adjacent brain section in competition with nicotine (300 |jM) (autoradiography scale bar: 0-1000 DLU/mm^2^); (K). [^18^F]nifene binding in DLU/mm^2^ to GM and WM of all CN TCX and FCX; (L). Average [^18^F]nifene binding in DLU/mm^2^ to GM and WM within CN TCX and FCX. Paired two-tailed parametric t-tests determined statistical significance between GM and WM (**p<0.01,***p<0.001).

### 3.4 Comparison of CN, DSAD and AD using nicotine effect

Figure 5 summarizes the extent of [^18^F]nifene binding within DSAD, AD, and CN in the two brain regions. When comparing between all groups within TCX (Figure 5A), FCX (Figure 5B), and all together (Figure 5C), the GM [^18^F]nifene binding was noticeably higher compared to the WM. The FCX exhibited marginally higher binding compared to TCX. Between the 3 groups no significant differences in GM binding as well as WM [^18^F]nifene binding was observed. There was also no significant difference in GM/WM [^18^F]nifene binding ratios (Figure 5D). Because WM has been previously reported to contain α4β2* nAChRs (Bieszczad et al. 2012; Mukherjee et al. 2018), the use of WM as a reference region may not be suitable. Variability in WM [^18^F]nifene binding in the 3 groups is also expected and therefore this would influence the GM/WM ratios. A more appropriate ratio comparison would therefore be the use of nonspecific binding measures as discussed below.

**Figure 5:**
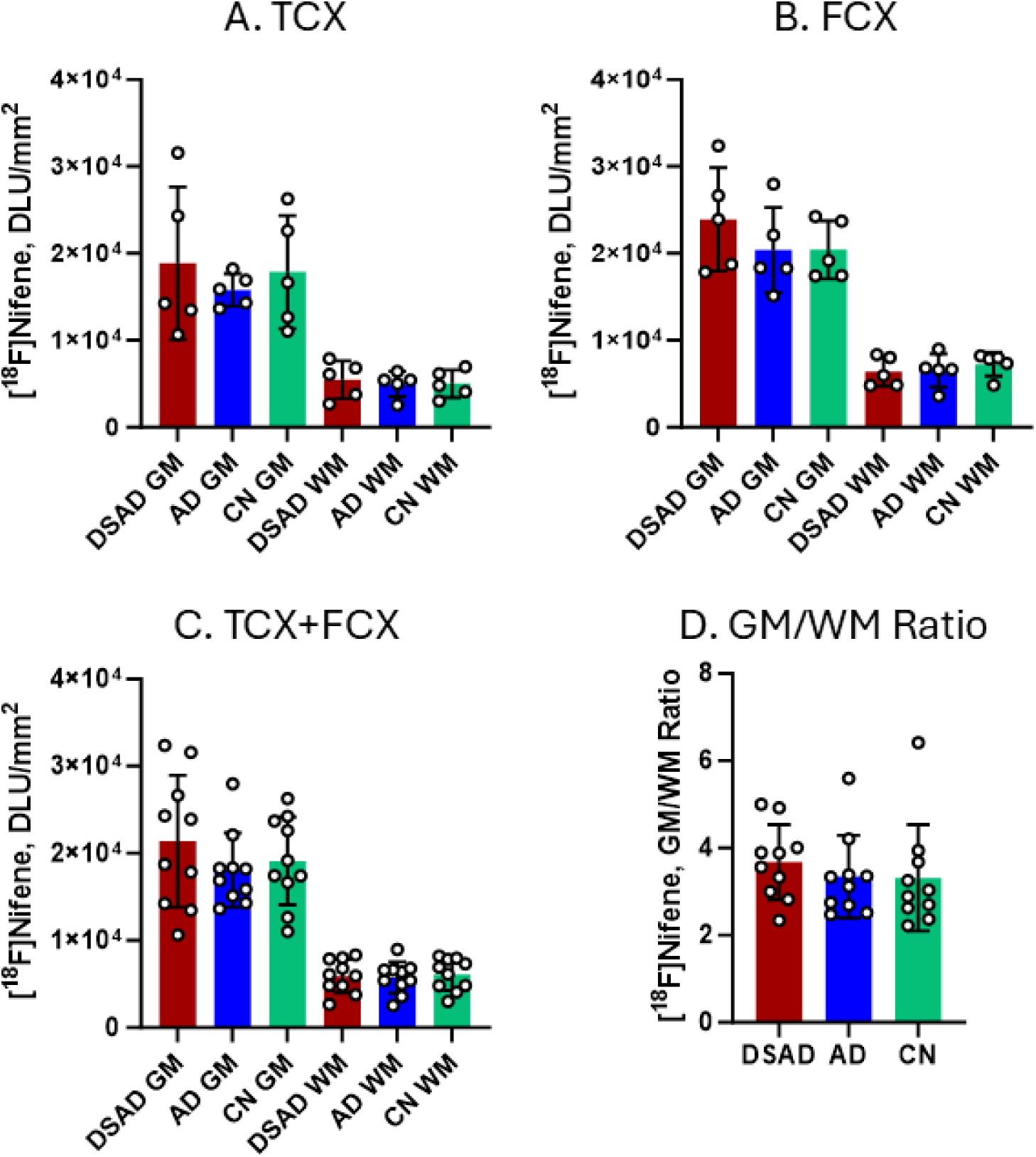
GM and WM comparisons all cases: [^18^F]nifene binding comparisons between DSAD, AD, and CN in (A). TCX (B). FCX (C). TCX and FCX (D). GM/WM ratios of [^18^F]nifene binding to TCX and FCX in all groups.

### 3.5 Relationship between [^18^F]Nifene, [^125^I]IBETA and [^18^F]MK-6240

Nicotine was able to displace [^18^F]nifene binding very efficiently from the GM and WM regions in the 3 groups (Figure 2J,3J,4F), consistent with our previous in vitro and in vivo competition studies of nicotine with [^18^F]nifene (Pichika et al. 2006; Campoy et al. 2021; Kant et al. 2011; McVea et al. ). The extent of displacement nicotine had on [^18^F]nifene was compared to total [^18^F]nifene binding to GM in adjacent slices (Figure 6). In TCX GM, nicotine displaced [^18^F]nifene binding by 87%, 89%, and 93% in DSAD, AD, and CN respectively (Figure 6A). In FCX GM of DSAD, AD, and CN, nicotine displaced [^18^F]nifene binding by 91%, 90%, and 95% respectively (Figure 6B). The ratios of GM binding to [^18^F]nifene+nicotine binding were the greatest in CN (TCX=14.6; FCX=19.4), while AD (TCX=10.4; FCX=10.6) and DSAD (TCX=7.8; FCX=11.7) were lower (Figure 6C). Average ratios of both the brain regions were CN=17, AD=10.5 and DSAD=9.74, suggesting a 38% decrease in AD and a 43% decrease in DSAD of [^18^F]nifene binding compared to CN (Figure 6D).

**Figure 6:**
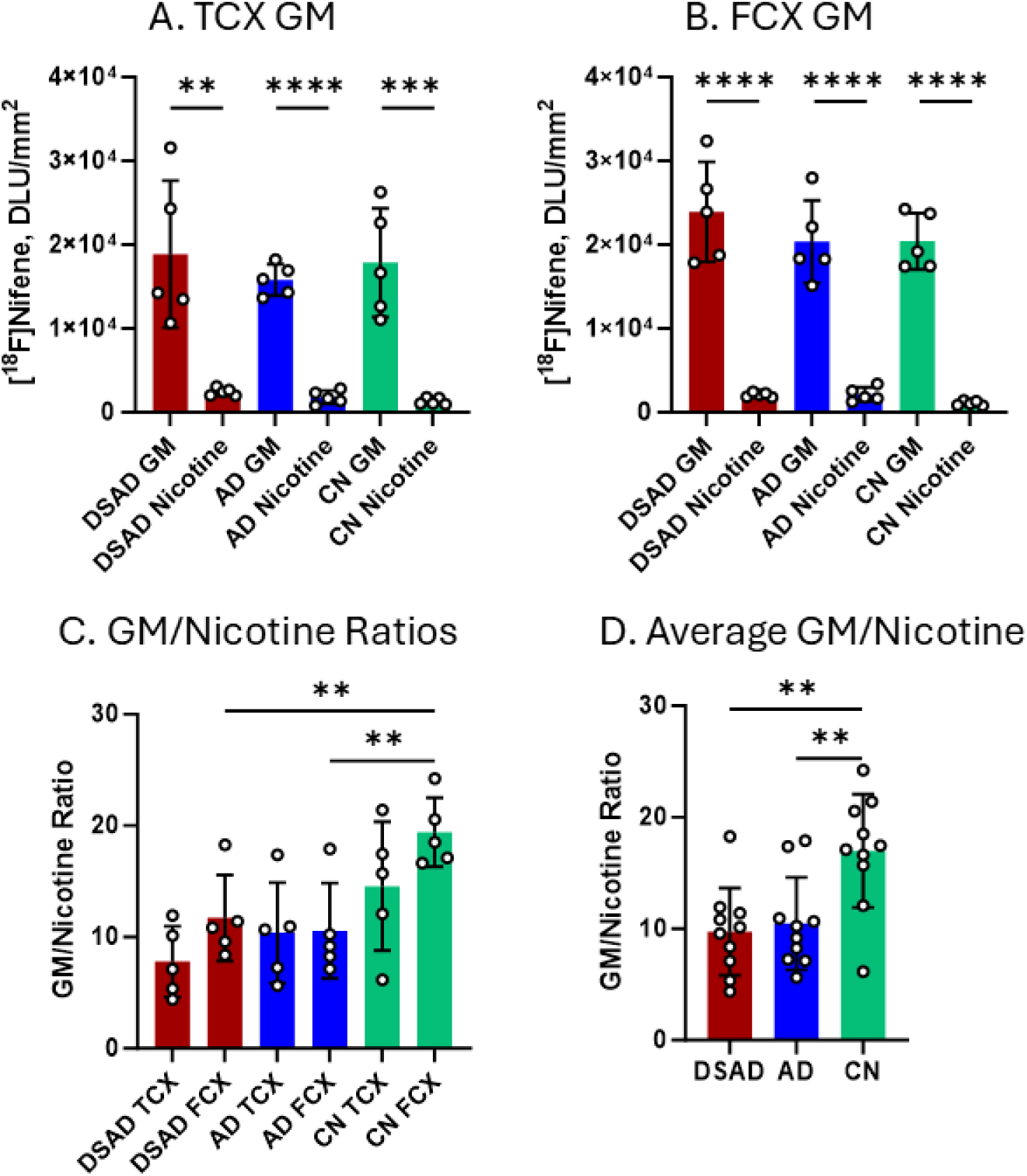
GM and nicotine competition comparisons: (A-B). [^18^F]nifene total binding in GM compared to [^18^F]nifene binding in competition with nicotine to TCX (A) and FCX (B) of all groups. Unpaired two-tailed parametric t-tests determined statistical significance between GM and nicotine competition (**p<0.01, ***p<0.001, ****p<0.0001); (C). Ratios of total binding to GM and [^18^F]nifene+nicotine binding in TCX and FCX of all groups. One-way ANOVA with Dunnett’s test determined statistical significance of CN compared to DSAD and AD (**p<0.01); (D). Average ratios of total binding to GM and [^18^F]nifene+nicotine binding. One-way ANOVA with Dunnett’s test determined statistical significance of CN compared to DSAD and AD (**p<0.01).

The effects of nicotine on WM [^18^F]nifene binding was evaluated separately. The extent of displacement nicotine had on [^18^F]nifene in the WM was lower than that observed for GM (Figure 7). In TCX WM, nicotine displaced [^18^F]nifene binding by 56%, 64%, and 74% in DSAD, AD, and CN respectively (Figure 6A). The nicotine-induced reductions seen in the WM were lower compared to GM, suggestive of some nonspecific [^18^F]nifene binding in WM. In FCX WM of DSAD, AD, and CN, nicotine displaced [^18^F]nifene binding by 67%, 67%, and 85% respectively (Figure 7B). The ratios of WM binding to [^18^F]nifene+nicotine binding were the greatest in CN (TCX=4.35; FCX=6.90), while AD (TCX=3.30; FCX=3.32) and DSAD (TCX=2.33; FCX=3.13) were lower (Figure 6C). Average WM ratios of both the brain regions were CN=5.62, AD=3.31 and DSAD=2.73, suggesting a 41% decrease in AD and a 51% decrease in DSAD of [^18^F]nifene binding compared to CN (Figure 7D).

**Figure 7:**
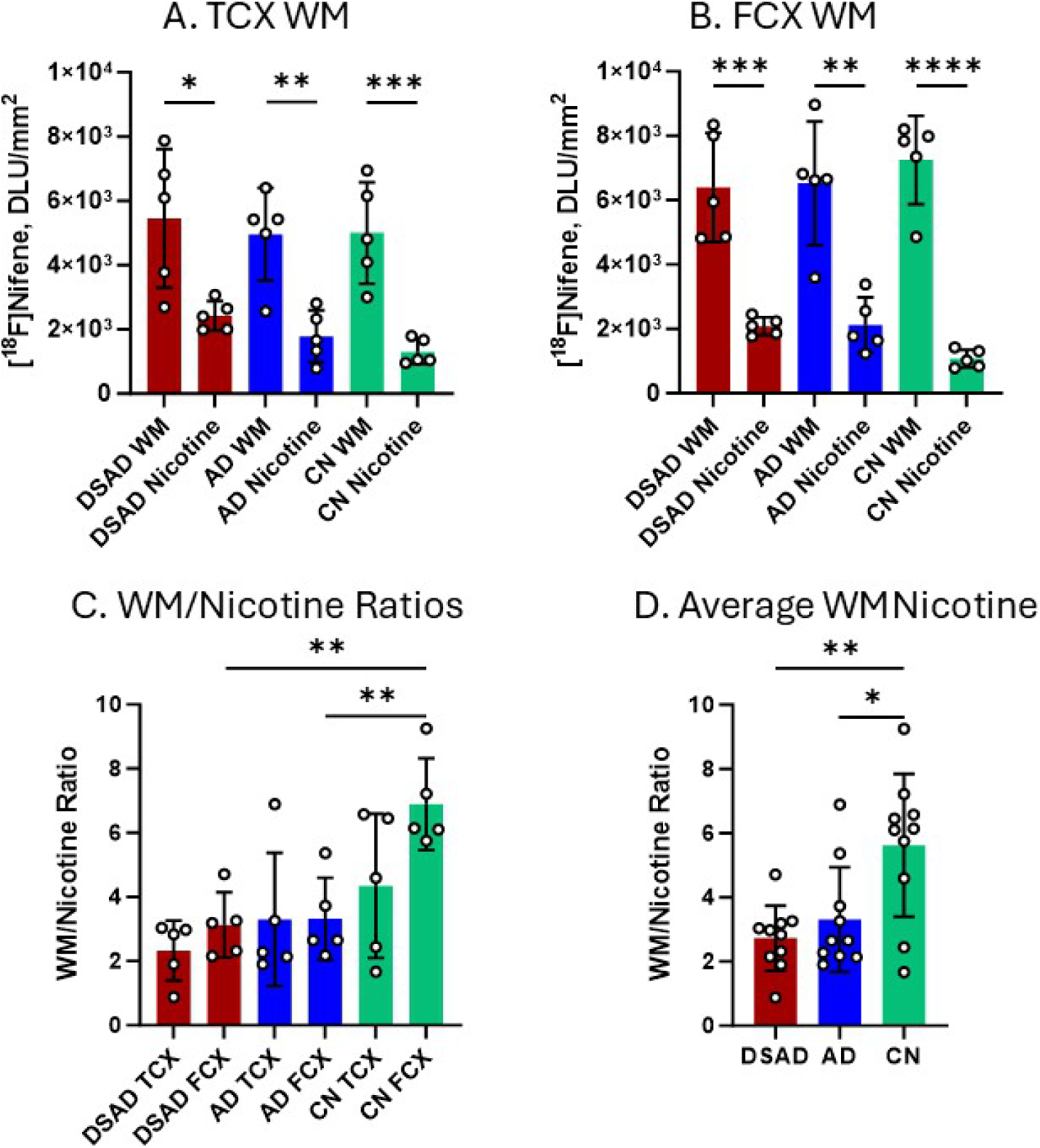
WM and nicotine competition comparisons: (A-B). [^18^F]nifene total binding in WM compared to [^18^F]nifene binding in competition with nicotine to TCX (A) and FCX (B) of all groups. Unpaired two-tailed parametric t-tests determined statistical significance between WM and nicotine competition (*p<0.05, **p<0.01, ***p<0.001); (C). Ratios of total binding to WM and [^18^F]nifene+nicotine binding in TCX and FCX of all groups. One-way ANOVA with Dunnett’s test determined statistical significance of CN compared to DSAD and AD (**p<0.01); (D). Average ratios of total binding to WM and [^18^F]nifene+nicotine binding. One-way ANOVA with Dunnett’s test determined statistical significance of CN compared to DSAD and AD (**p<0.01).

Thus, both the GM and WM in AD and DSAD exhibited significant decreased [^18^F]nifene binding in different brain regions. The decrease in DSAD was greater than in AD.

### 3.6 Relationship between [^18^F]Nifene, [^125^I]IBETA and [^18^F]MK-6240

The characteristics of α4β2* nAChR in DSAD can be further characterized in correlations with Aβ and tau (Figure 8). Adjacent slices of the same cases were used for [^125^I]IBETA and [^18^F]MK-6240 binding to Aβ and tau respectively (Biju et al. 2026; Karim et al. 2026). A strong positive correlation between Aβ ([^125^I]IBETA) and α4β2* nAChR ([^18^F]nifene) was observed in DSAD FCX and TCX (Figure 8A), suggesting an association of Aβ-α4β2* nAChR. However, in AD such a strong Aβ-α4β2* nAChR correlation was not seen and is consistent with our previous work in the AD hippocampus-subiculum (Karim et al. 2025).

**Figure 8:**
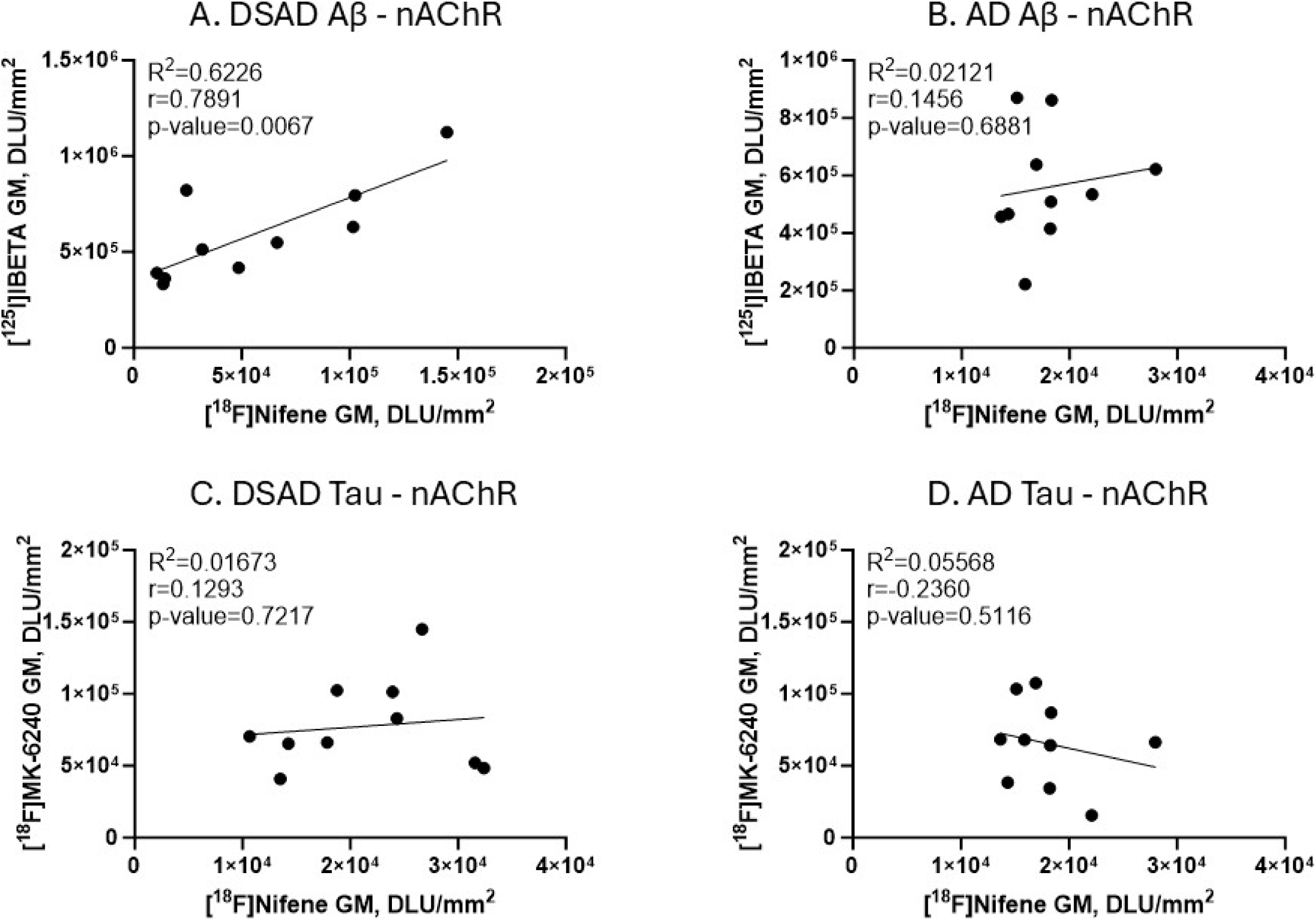
[^18^F]Nifene, [^125^I]IBETA and [^18^F]MK-6240 binding in DSAD and AD: (A). Correlation between Aβ and α4β2* nAChR in DSAD (R^2^=0.6226, Pearson’s r=0.7891, p-value=0.0067); (B). Correlation between Aβ and α4β2* nAChR in AD (R^2^=0.02121, Pearson’s r=0.1456, p-value=0.6881); (C). Correlation between tau and α4β2* nAChR in DSAD (R^2^=0.01673, Pearson’s r=0.1293, pvalue= 0.7217); (D). Correlation between tau and α4β2* nAChR in AD (R^2^=0.05568, Pearson’s r=0.2360, p-value=0.5116).

In the case of correlations between tau ([_18_F]MK-6240) and α4β2* nAChR, there was weak correlation in the case of both DSAD and AD. In the case of AD, a weak negative trend was observed for tau-α4β2* nAChR, which was similar to our observations in the AD hippocampus-subiculum (Karim et al. 2025).

### 3.7 Age correlation of [^18^F]Nifene binding

Figure 9 shows a correlation between age and [^18^F]nifene GM and WM ratios in the three groups, DSAD, AD, and CN. Ratio of total binding/nonspecific binding measured in the presence of nicotine was plotted for TCX and FCX., separately for GM and WM. The DSAD cases were the youngest while the CN cases were the oldest, with some overlapping with AD cases. The CN cases exhibited the highest GM ratios in both the TCX and FCX (>15) and trended upwards with age. Both DSAD and AD GM ratios were similar (210), and thus lower compared to CN. As expected, [^18^F]nifene binding ratios in the WM were lower compared to the GM. However, the ratios followed similar trends as the GM, with WM ratios in the CN cases being higher than AD and DSAD cases. There was a positive correlation with age in TCX and FCX in the combined plots, but more precise aging effect in each group will have to be obtained by increasing the number of cases in order to separate aging and disease effects. The small number of cases in the present study did not allow a precise male-female comparison.

**Figure 9:**
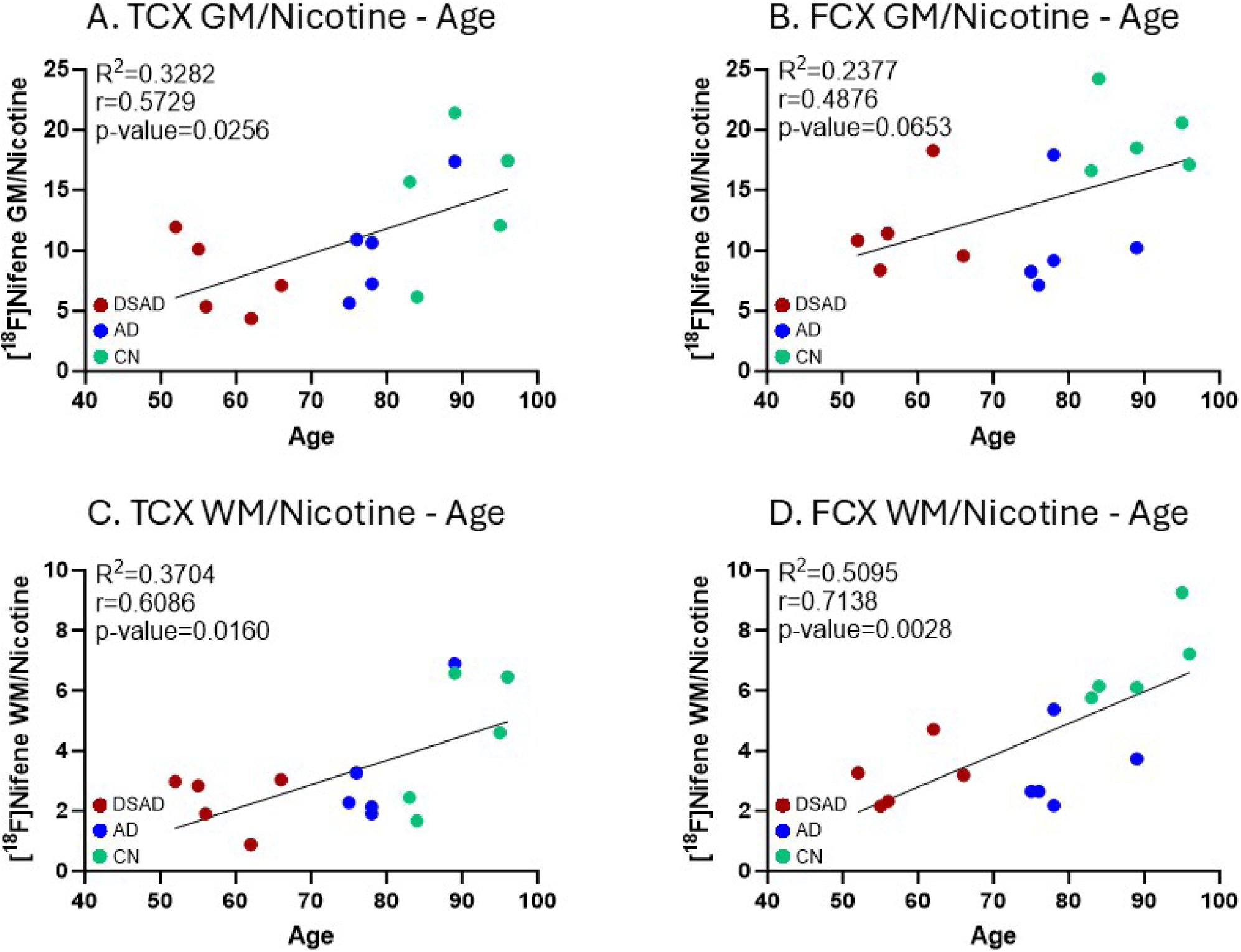
[^18^F]Nifene correlated with age in all cases: (A). Correlation between age and [^18^F]nifene GM/nicotine ratios in TCX of DSAD, AD, and CN (R^2^=0.3282, Pearson’s r=0.5729, p-value=0.0256); (B). Correlation between age and [^18^F]nifene GM/nicotine ratios in FCX of DSAD, AD, and CN (R^2^=0.2377, Pearson’s r=0.4876, p-value=0.0653); (C). Correlation between age and [^18^F]nifene WM/nicotine ratios in TCX of DSAD, AD, and CN (R^2^=0.3704, Pearson’s r=0.6086, p-value=0.0160); (D). Correlation between age and [^18^F]nifene WM/nicotine ratios in FCX of DSAD, AD, and CN (R^2^=0.5095, Pearson’s r=0.7138, p-value=0.0028).

## 4. DISCUSSION

As brain regions deeply involved in AD and DSAD progression, investigating the role of α4β2* nAChRs in FCX and TCX relative to known, prevalent biomarkers can reveal more details in disease pathology and severity. This study revealed selective [^18^F]nifene binding in the FCX and TCX of DSAD cases, comparable to AD and CN. With ratios of GM or WM to nicotine presence, DSAD and AD were significantly less than CN. The most noteworthy correlations were between [^18^F]nifene and [^125^I]IBETA in DSAD GM and [^18^F]nifene GM/WM ratios with age of all groups.

As an acetylcholinesterase inhibitor used conventionally in AD, donepezil has been investigated in DS with AD-associated cognitive impairment or dementia (Lott et al. 2002; Prasher et al. 2002; Prasher et al. 2003). Clinical studies with only DS patients have yet to determine a consensus on the clinical benefit of donepezil (Johnson et al. 2003; Spiridigliozzi et al. 2007; Kishnani et al. 2010; Kondoh et al. 2011). The improvement of AD-related symptoms in treatment with donepezil is evident while results are mixed in only DS, suggesting that its effect may be specific to AD pathology. These efforts to validate AD therapeutics in DSAD can inspire the application of detecting AD pathology in DSAD, including cholinergic deficits. The Ts65Dn model of DS reflects cholinergic system deterioration in which decreased cholinergic fibers led to changes associated with atrophy of basal forebrain cholinergic neuron (BFCN) (Isacson et al. 2002; Chen et al. 2009). Many more studies in Ts65Dn mice and humans with DSAD demonstrate a deficiency of BFCNs and reduction in basal forebrain volume to highlight impaired cholinergic activity in DS (Granholm et al. 2000; Powers et al. 2016; Aranha et al. 2023). The connectivity of the cholinergic basal forebrain receives brainstem and limbic input to influence cholinergic innervation in cortical and neocortical regions, also affected by the loss of cholinergic input in neurodegenerative diseases (Martinez et al. 2021; Cykowski & Masdeu 2025). Loss of input downregulates the expression of cholinergic receptors. There is an overall reduction of α4β2* nAChRs in AD compared to CN while findings are inconclusive for α7 nAChRs, another abundant subtype (Sabri et al. 2018; Coughlin et al. 2020; Karim et al. 2025; Ngo et al. 2025). Amyloidogenic peptides Aβ1-42 are pathologically overproduced in DS, therefore binding to α7 and α4β2* nAChRs in greater extents which induces cytotoxicity and amyloid plaque formation that underlie cognitive deficits (Wang et al. 2000a; Wang et al. 2000b; Deutsch et al. 2014; Keihan Falsafi et al. 2016). The binding affinity of Aβ1-42 to α7 nAChRs is higher, requiring greater concentrations of Aβ1-42 to bind to α4β2 nAChRs. Maximum plaque burden is likely attained in all the DSAD and AD cases with high plaque stages (Table 1) so the effects on α4β2* nAChRs may be indicated. The positive correlation between Aβ ([^125^I]IBETA) and α4β2* nAChR ([^18^F]nifene) was observed in DSAD (Figure 8A), may suggest such an association of Aβ-α4β2* nAChR.

Despite the traditional concept of GM abnormalities driving AD earlier than WM, abnormalities in WM may also play contributing roles (Ge et al. 2002). Significant reductions in WM [^18^F]nifene binding in DSAD and AD compared to CN suggests loss of WM microstructural integrity. Such losses may account for clinical symptoms of mild cognitive impairment, associated with cognitive decline (Vernooij et al. 2009; Shafer et al. 2022). Despite atrophy, basal forebrain GM volume positively relates with the reconfiguration of WM networks that support residual memory (Ray et al. 2015). With the association between WM integrity and cognition established, the microstructural properties of WM can indicate cognitive decline in neurodegenerative diseases. The frontal tracts particularly suffer from lower WM integrity in DS compared to CN, involving connectivity differences that result in praxis deficits (Powell et al. 2014). In DS, Aβ burden was associated with widespread longitudinal WM microstructural changes detected in diffusion tensor imaging (LeMerise et al. 2025). Significant relationships with Aβ and tau burden suggest a link between WM disruptions and multiple AD-specific mechanisms (Bazydlo et al. 2022; Edwards et al. 2024). WM degeneration in DSAD precedes AD symptom onset and cognitive decline, posing as a potential diagnostic avenue with fluid and molecular biomarkers (Morcillo-Nieto et al. 2024; Silva et al. 2026). As WM degeneration worsens with AD progression, the WM microstructure and connectivity to various brain regions are sensitive to cognitive decline starting early in DSAD. Further investigations of WM abnormalities in the context of multiple molecular pathologies can be valuable.

The overexpression of APP in DS leads to nerve growth factor transport and cholinergic neurodegeneration (Salehi et al. 2006). Although independent of Aβ, APP is required for the overactivated rab5-mediated pathway and endosome dysfunction in DS and AD (Jiang et al. 2009). In this study with only Aβ-positive cases, a significant positive correlation between Aβ and α4β2* nAChRs was observed. The abnormally high accumulation of soluble Aβ may interfere with nAChRs and ultimately compromise the cholinergic system in DS (Deutsch et al. 2003). Cholinergic overactivity in early stages of DS may contribute to cognitive impairment while later stages exhibit a widespread deficiency (Lisgaras & Scharfman 2026). This dynamic shift in cholinergic signaling can potentially be detected using [^18^F]nifene binding to α4β2* nAChRs.

The preliminary findings in this study may have the following limitations. All the DSAD and AD cases were Braak stage VI with substantial plaque burden. Although this consistency allows for implications of greater disease severity, the development in proteinopathy is unable to be accurately estimated. The small number of cases in each group limits the broader application of these findings so greater group sizes are necessary in subsequent studies. Increased group size can minimize the effect of inter-case variability, which these results were unaffected by. Additionally, the ages of CN were noticeably greater than DSAD and AD so age correlations may not accurately account for younger CN. Older CN exhibits the greatest tau and plaque burden from aging, granting closer comparisons to the extents achieved by disease severity albeit the advantages of age-matched cases. Despite limitations, this study provides sufficient validation of [^18^F]nifene and its role as a supportive radiotracer in elucidating DSAD pathophysiology and diagnosis. In summary, [^18^F]nifene binding to α4β2* nAChRs has the capability to accentuate unique cholinergic effects in DSAD FCX and TCX similarly to AD and CN.

## ACKNOWLEDGEMENTS

Financial support for the project was provided by NIH AG 029479 and AG 077700. We thank Jeffrey Kim, Pathology and Laboratory Medicine, University of California-Irvine for immunostaining of brain sections.

## Author Contributions

All authors had access to all the data and assess the accuracy of the data analysis. Conceptualization, J.M.; methodology, F.K., C.L., N.P.C-J.; software, F.K., N.P.C-J..; validation and analysis, F.K., C.L. J.M.; investigation, J.M..; resources, E.H.; writing—original draft preparation, F.K., J.M.; writing—review and editing, FK, E.H., JM; supervision, J.M..; project administration, J.M..; funding acquisition, J.M.

## Data Sharing

The data that support the findings of this study are available from the corresponding author upon reasonable request.

